# Nutri rich cattle feed (NuCa feed) supplement with improved fiber content free of anti-nutritional factor Phytic acid

**DOI:** 10.1101/2020.08.28.271718

**Authors:** Shivakiran Makam, Harish Babu Kolla, Daya Mouni Golla, Ananya Payal, Meghanath Somarowthu

## Abstract

Most of the commercial ruminant feeds comprise Corn meal, Groundnut cake, Maize etc., which contain anti-nutritional factors called Phytates. They have high binding affinity to cationic minerals. This results in decrease in activity of the cattle by making minerals unavailable for absorption in the intestine. Many of the feed industries add Phytase enzyme that degrades the Phytic acid. Addition of Phytase enzyme is very complex method and increases the cost of the feed. If Phytase is not added, Phytic acid enters the ecosystem through the dung and due to microbial degradation, the Phytic acid gets converted into Phosphates. Phosphate abundance in water bodies leads to Eutrophication, which is a serious ecological issue. And also, in many parts of the country, post-harvest Paddy straw is burnt unused. This intentional stubble burning causes air pollution. We combined paddy straw and Azolla along with jaggery to prepare animal feed supplement. The feed pellets were evaluated for their nutritional composition and efficacy. The efficacy of the developed feed was evaluated by feeding it to milch cows and compared with regular feed for a period of 15 days. The Azolla-Hay feed improved the milk yield and also quality of the milk in comparison to the control group. This study revealed that Azolla in combination with paddy straw powder can be low-cost feed alternative especially during the lean period where the availability of feed is scarce.

## Introduction

Livestock industry in India contributes significantly towards the Gross Domestic product (GDP). It ensures welfare of the rural population. Most of the agricultural families depend on livestock maintenances for their livelihood. And also, this sector provides supplementary employment to the marginal farmers in rural areas. Thus, Livestock emerging as an important sector, leverage the rural economy (1). Although livestock sector has greater growth potential, the further growth of the sector mainly depends upon the availability of fodder. The major challenge is shortage of fodder during summer and drought conditions. The feed comprised of Groundnut cake, Soybean, Corn meal etc. given as nutritional supplements during lean periods of drought or summer. Phytic acid (Phytate) is an anti-nutritional factor is present in the feed and resembles Glucose molecule in structure (presence of phosphate groups in hydroxyl positions of glucose). It is very essential in holding the minerals which later released during germination (2, 3, 4, 5, 6). These anti-nutritional Phytates have high affinity towards the Minerals such as Calcium, Magnesium, Iron, Copper, and Zinc ^5^. This chelation results in precipitation, making minerals unavailable for absorption in the gut. Hence, activity of the cattle decreases (6). To overcome this problem, many of the feed industries add immobilized enzyme Phytase. Phytase is a type of Phosphatse enzyme that degrades phosphate groups in Phytic acid molecules in intestine of Bovines ^7^. However, enzyme addition increases the production cost of the feed. If Phytase is not added, Phytic acid comes out to ecosystem through the dung and gets degraded into Phosphates by microbial action. Increased Phosphate in water bodies leads to Eutrophication, which is a serious ecological issue (8, 9, 10). Since ancient times Paddy straw is used as conventional silage to the cattle and other animals. It increases the fat content in the milk and promotes milk quality(11). Similar studies with Azolla show that the animals produce more milk by 10-12 % when fed them with Azolla(12). Azolla is a free floating Pteridophyte that belong to family Salvinaceae in Plant kingdom. It contains blue-green algae, an endosymbiont that fixes atmospheric nitrogen and produce a variety of proteins (13, 14). Azolla multiplies by splitting or budding (15). It produces the large amount of biomass (1000 MT/Hectare/year) and very rich in proteins (25-30 MT protein/hectare/year) (16). Besides, the hay and Azolla are free from Phytates. The hay and Azolla are easy to procure and produce. In many parts of the country, hay after the crop is burned unused which leads to air pollution. Hence in the current study, we have attempted to prepare low-cost nutritional feed to cattle and made available throughout the year. Such low-cost feed combinations will be of great help to countries that mainly depend on agriculture especially in tropics. We have also studied its nutritional properties and effects on milk yield by feeding them to milch cows for 15 days.

## Materials and Methods

### Materials

Azolla culture, Nutrimin and super Phosphate powder were procured from Sangam dairy, Vadlamudi, Guntur District, AP, India. Paddy straw was collected from nearby paddy fields in the village. Paddy straw chopping machine and pulverizer were purchased from 6S Mechatronics, Coimbatore, Tamil Nadu, India. Pellet making machine was purchased from Vishwatech Fabrications, Nagpur, Maharashtra, India.

The materials for preparing Azolla beds-bamboo poles, HDPE tarpaulin sheets (350 GSM) of 3×6 feet in size, shade nets and ropes were procured from local market. We preferred 350 GSM tarpaulin sheets because they would last for at least one year in direct sunlight. The perforated nets or mosquito net sheets for Azolla dehydration were procured from local market. The iron stands for setting up of dehydrating Azolla were fabricated at the University in-house workshop.

### Azolla cultivation and Hay powder preparation

#### Preparation of paddy hay powder

Paddy hay from the fresh paddy harvest was collected and chopped to1-2 inches in the chopping machine and then powdered in Pulverizer.

#### Preparation of Azolla Beds and Azolla cultivation

The pictorial representation of preparation of Azolla beds is shown in figure 2. The bamboo poles were cut to two sizes-6 ½ feet and 3 ½ feet in length. For each bed, two numbers of former and eight numbers of latter were used. The tarpaulin was spread and tied to the bamboo poles are shown in the picture. About 20 kgs of sieved red soil was uniformly spreaded over the sheet. Fresh cow/buffalo dung is dried in sunlight till the moisture was eliminated completely. Then, slurry was prepared with dried dung of 4 kgs mixed in 10 liters of water and poured into the Azolla cultivation bed uniformly. Then the bed was filled with water up to 9 inches height. About 0.5 kg of a pure Azolla mother culture was inoculated into the bed uniformly. Within a week, Azolla had spread completely in the bed and appeared like a thick mat (17, 18, 19). The physical and chemical requirement for optimum growth of Azolla was maintained as shown in table 1. A mixture of 20 gm of Nutrimin (table 2), 50 gm of super phosphate and about 1 kg of dried dung was added once in 07 days to maintain rapid multiplication of the Azolla and 500 gm of daily yield (20). About 5 kg of bed soil was replaced with fresh soil once in 30 days to avoid nitrogen build-up and to prevent nutrient deficiency. 25 to 30 percent of the water also replaced with fresh water for every 10 days to prevent nitrogen build up in the bed.

**Table 1.**
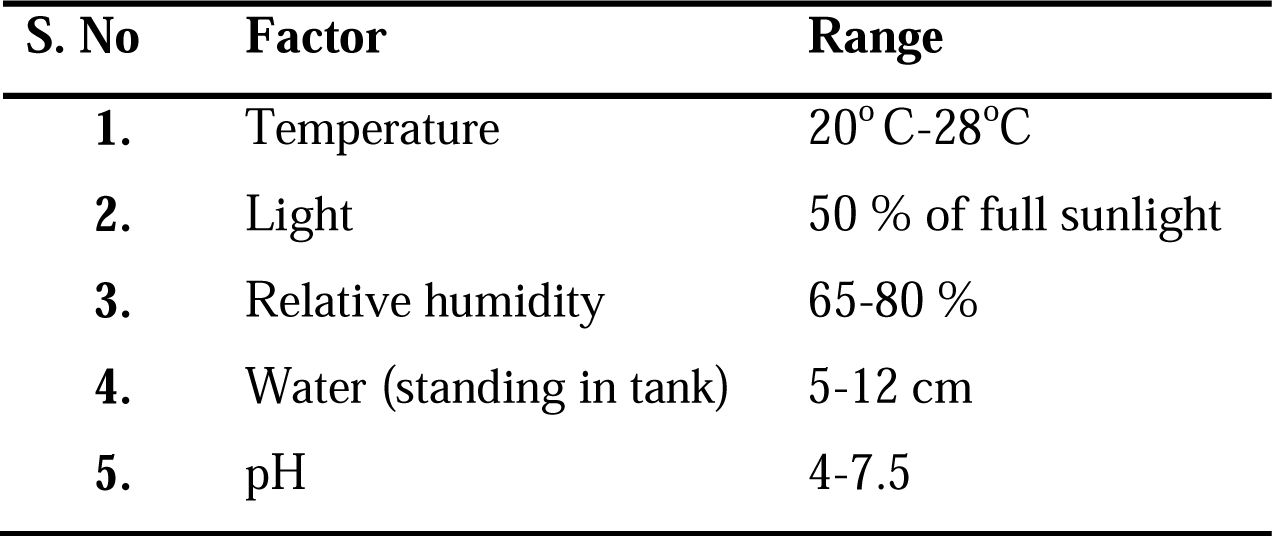
Factors required for the growth of Azolla.

**Table 2.** Mineral composition of Mineral mixture used for Azolla growth (as mentioned on the product pack).

| S. No | Components | Composition<br>(for 100 gms) |
| --- | --- | --- |
| 1. | Vitamin A | 75,000 i.u |
| 2. | Vitamin D<sub>3</sub> | 7,500 i.u |
| 3. | Vitamin E | 35 mg |
| 4. | Calcium | 25 g |
| 5. | Phosphorous | 12 g |
| 6. | Magnesium | 2 g |
| 7. | Zinc | 675 mg |
| 8. | Manganese | 432 mg |
| 9. | Copper | 108 mg |
| 10. | Iodine | 43 mg |
| 11. | Cobalt | 108 mg |
| 12. | Selenium | 3 mg |
| 13. | Chromium | 2 mg |
| 14. | Yeast | 4 g |

**Figure 1.**
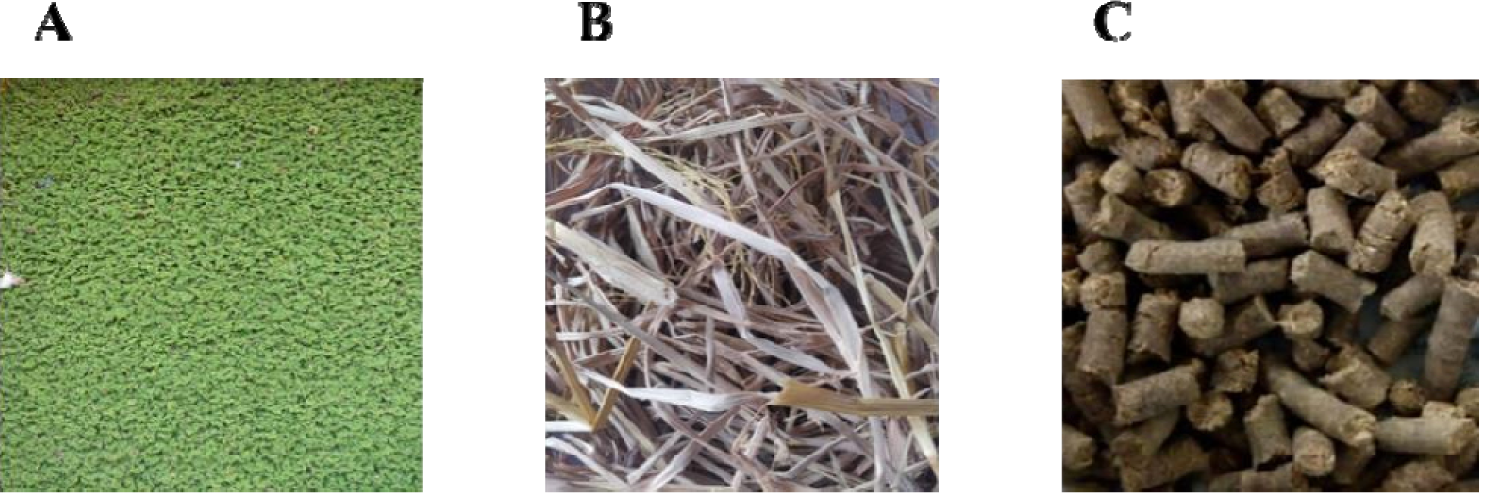
Feed preparation by the combination of Azolla and Paddy straw. **A.** Azolla cultivated in 6ft×3ft×1ft rectangular metal stand. **B.** Paddy straw collected from fields. **C.** Feed pellets prepared by the combination of Azolla and Paddy straw by pellet technology.

**Figure 2:**
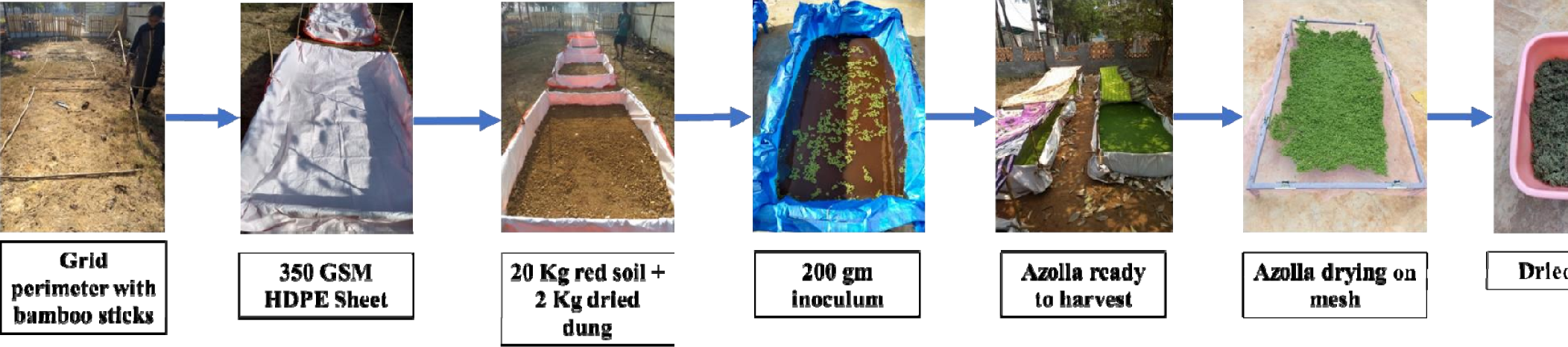
Preparation of Azolla Grids for cultivation & drying.

### Feed Preparation and Nutritional Analysis

Azolla multiplied rapidly within 10 - 15 days. From then, 500 - 600 gm of Azolla was harvested daily. Harvesting was done every day from the 15^th^day onwards using plastic sieve. Harvested Azolla was washed with fresh water to get rid of the cow dung smell. Fresh Azolla collected was sundried for 4-6 hours to remove the water content in Azolla. Dried Azolla was passed through the pulverizer for fine powdering. Finely powdered hay and Azolla were mixed with the jaggery as binder in an 8:1:1 ratio (8 parts of Hay powder, 1 part of Azolla and 1 part of Jaggery by weight). Then, the powdered form of feed was made into pellets in the Pelletizer. Proximate analysis of the prepared feed concentrate was performed. The proximate analysis was outsourced to Eurofins Nutritional Analysis Center, Eurofins, Bangalore, Karnataka, India. Proximate analysis revealed the nutritional content of the feed. The prepared feed pellets were fed to cows to check the milk yield per each cow.

### Animal trials and evaluation of NuCa Feed on cows

NuCa Feed was assessed for its potential on cows. For this, 4 milch cows, Jersey crossbreed were considered. They were grouped as control (C1 & C2) and the test (T1 & T2) with two animals in each group. Prior to the experimentation, health status of the animals was checked, and they were considered for trials. The control group was fed with 2 Kgs of regularly available feed concentrate along with the fodder. Then the test group received the regular fodder and additionally 2 kgs of NuCa Feed prepared in this study. This regime was continued for 15 days. During starting day (Zeroth day), seventh day and fifteenth days of the experiment, the following parameters were measured-(i) milk yield per day, (ii) density of milk & (iii) lactometer reading.

## Results and Discussion

### Azolla cultivation and Hay powder preparation

After the inoculation of Azolla in the beds, Azolla had grown completely in one week occupying the total surface of the water. Each bed had yield 400-600 gm of Azolla every day for about a week. Then, the yield was found to decrease. Adding additional amount of super phosphate and dried cow dung improved the yield. After drying, about 70-100 gm of Azolla was obtained from one kg of fresh Azolla. There was variation in the yield of Azolla per bed and also in the yield of dried Azolla. Meanwhile, pulverized paddy straw was prepared from freshly harvested paddy fields. The paddy straw, Azolla and jaggery were mixed in 8:1:1 ratio. The NuCa Feed pellets were prepared in Pelletizer. About 25 kg feed pellets were prepared in one hour. The NuCa Feed pellets were packed in water-proof covers for further use.

### Proximate nutritional analysis of NuCa feed

Results of proximate analysis of the NuCa Feed, azolla powder and the hay powder are represented in table 3. Azolla was rich in proteins (15.35 gm); and minerals like Iron (461.71 mg), Calcium (8348.01 mg) and Potassium (1281.30 mg), whereas hay powder was rich in fiber (73.63 mg), Energy (322.376 kcal) and total Carbohydrates(73.63 gm).

**Table 3.** Proximate Nutritional analysis of Azolla powder and hay powder.

| S.No. | Components | Azolla Powder<br>(per 100 g) | Hay Powder<br>(per 100 g) | NuCa Feed<br>(per 100 g) |
| --- | --- | --- | --- | --- |
| 1. | Energy(kcal) | 228.62 | 322.37 | 324.13 |
| 2. | Total carbohydrates(g) | 33.66 | 70.21 | 66.48 |
| 3. | Dietary fiber on dry basis(g) | 32.57 | 73.63 | 50.68 |
| 4. | Ash(g) | 38.01 | 13.23 | 15.15 |
| 5. | Total fat(g) | 3.62 | 2.21 | 3.93 |
| 6. | Protein(g) | 15.35 | 5.41 | 5.71 |
| 7. | Iron(mg) | 461.71 | 88.51 | 247.05 |
| 8. | Calcium(mg) | 8348.01 | 413.04 | 2150.57 |
| 9. | Potassium(mg) | 1281.30 | 1409.16 | 1120.78 |

In table 4, test feed is compared with the two commercial feeds A and B in terms of components, fate of phytates in them and advantages of NuCa feed over the commercial feeds. Feed A is composed of Groundnut cake, Corn, etc. whereas feed B comprises Maize, Soybean and Sunflower seeds. Phytates were present in both A and B since they are made of Cornmeal, Groundnut cake, Soya, Maize, etc. The test feed comprises Hay powder, Azolla powder and Jaggery was free of Phytates.

**Table 4.** Comparison of prepared feed with two commercial feeds A and B.

|  | Feed A | Feed B | Test feed (Nuca feed) |
| --- | --- | --- | --- |
| Components | Groundnut cake,<br>Corn, etc. | Maize, Soybean,<br>Sunflower seeds. | Paddy straw powder<br>(Hay powder), Azolla<br>and Jaggery. |
| Phytates | Present | Present | Absent |
| Enzymes added | Immobilized Phytase | Immobilized Phytase | No Enzymes are added |
| Type | Powder | Powder | Pellets |
| Production time | Requires more time to produce as it depends on the availability of food crops | Requires more time to produce as it depends on the availability of food crops | Very fast. Azolla has faster growth. |
| Agricultural land occupancy | Take away cultivation land | Take away cultivation land | Doesn't require any agricultural land |
| Accessibility | Not easily accessible | Not easily accessible | Affordable and accessible to farmers. |
| Transportation loss | Powder can be lost during transportation | Powder can be lost during transportation | Pellets can be easily packed and transported |

### Animal trials and efficacy of NuCa feed

Four milch cows were grouped into control and test group with 2 cows in each group. C1 and C2 were control that were fed with 2 kgs of regular feed along with the fodder. And 2 kgs of test feed with regular fodder was given to T1 and T2 cows. Feed trial experimentation was done for 15 days. Milk yield and Lactometer reading of each cow are shown in table 5 and 6. On a whole, in the test group T1 and T2 yielded 0.5 and 1.6 liters more than 0^th^ day, respectively by the end of the study, whereas C1 and C2 cows did not show any increase in the milk yield and showed poor milk yiled. There was increase in milk by 6.66% and 26.6% in T1 and T2 respectively.

**Table 5.** Milk yield of Each cow in Animal trails.

| Milk Yield (in Litres per day) |  |  |  |  |
| --- | --- | --- | --- | --- |
| Cow | 0<sup>th</sup>Day | 7<sup>th</sup>Day | 15<sup>th</sup>Day | Average milk yield (in Litres) |
| Control Cow C1 | 8.4 | 8.0 | 8.0 | $8.13 \pm 0.23$ |
| Control Cow C2 | 4.4 | 4.6 | 4.4 | $4.46 \pm 0.11$ |
| Test Cow T1 | 7.5 | 7.4 | 8.0 | $7.63 \pm 0.32$ |
| Test Cow T2 | 6.0 | 7.0 | 7.6 | $6.86 \pm 0.80$ |

**Table 6.**
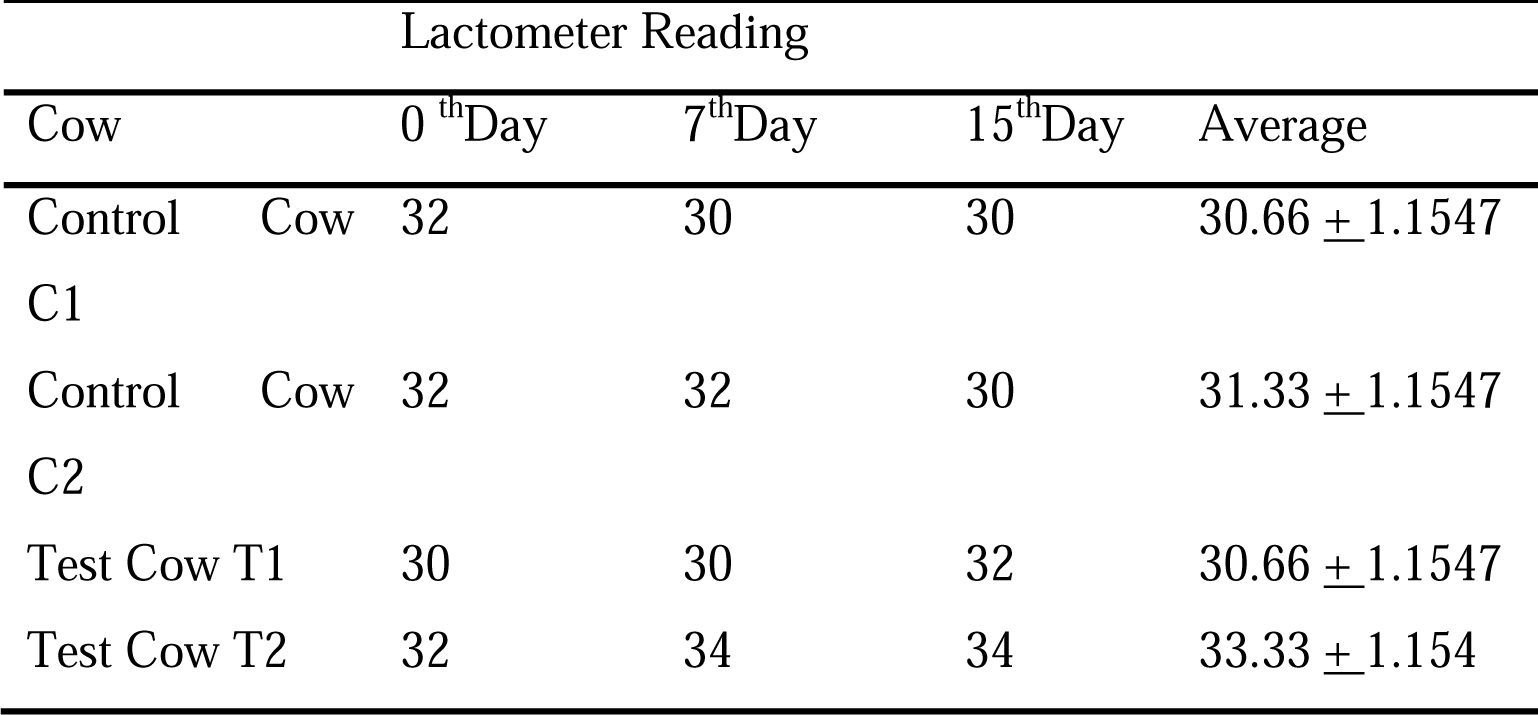
lactometer reading of milk.

Based on the Lactometer reading from table 6, density of milk were derived using the following equations:

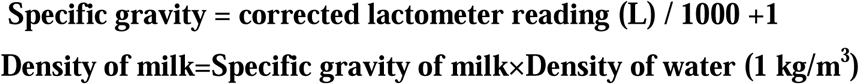

Density of milk was represented in table 7.

**Table 7.** Density of the milk based on lactometer reading.

| Cow | 0<sup>th</sup>Day | 7<sup>th</sup>Day | 15<sup>th</sup>Day | Average density<br>(kg/m<sup>3</sup>) |
| --- | --- | --- | --- | --- |
| Control Cow C1 | 1.032 kg/m<sup>3</sup> | 1.030 kg/m<sup>3</sup> | 1.032 kg/m<sup>3</sup> | 1.03133 ± 0.001154 |
| Control Cow C2 | 1.032 kg/m<sup>3</sup> | 1.032 kg/m<sup>3</sup> | 1.030 kg/m<sup>3</sup> | 1.03133 ± 0.001154 |
| Test Cow T1 | 1.030 kg/m<sup>3</sup> | 1.030 kg/m<sup>3</sup> | 1.032 kg/m<sup>3</sup> | 1.03066 ± 0.001154 |
| Test Cow T2 | 1.032 kg/m<sup>3</sup> | 1.034 kg/m<sup>3</sup> | 1.034 kg/m<sup>3</sup> | 1.03333 ± 0.0001154 |

In this study, the feed is prepared in a balanced ration (8:1:1), and also it is rich in protein and energy. Fiber content is high in this feed. High fiber helps in the normal functioning of gut in ruminants. The feed developed in the study comprises of Azolla, Hay powder and Jaggery. It is free of anti-nutritional factor Phytic acid. Since we used pellet technology, pellets has high digestibility and they are masticated by ruminant animals to extract more nutrients. Compared to regular format of feeding the animals with fodder, this pellet technology helps in dissolving and segregating the nutrients in the feed. Majority of the feed products available in the market exist in powder form. Instead we used pellet technology to make pellets. The Azolla-Hay pellets developed were fed to cows for 15 days and feeding behavior/affinity of the cows towards the feed is analyzed; milk yield, density and lactometer reading of the milk among the cows fed with the Azolla-Hay pellets in comparison with the regular commercial feed supplements was noted. This study has observed that the cows relished on the test feed and they were not averse or rejecting the feed. All the 15 days, the cows relished on the test pellets. There was no physical dullness or lethargy in the cows that were fed with test feed. The tables 3 and 5 above has shown that the pellet feed is rich in nutrients and improved the milk yield. The study proved that the test pellets could be the low cost alternative feed supplement for cows. Moreover, this study also shows that the production of Azolla is cheap and effective and the cost would be much lesser than the commercial feed supplements which are also having Phytate molecules that cause ecological effects. And also the feed developed in this study could be an effective alternative for stubble burning.

Feed shortage has become one of the major challenges to livestock sector. And also many of the commercially available feeds comprise of Corn, Groundnut, Soya, and Maize etc. These crops are also the food crops for humans and take away cultivation land widely. Another drawback of commercial feeds is that they comprise Phytic acid, an anti-nutritional factor. Phytic acid chelates with cationic minerals and reduces the absorption of minerals in the intestine. Many of the feed industries add enzyme Phytase. Addition of enzymes to feed is a very complex and costly procedure. It increases the production cost of the feed. If it is not done so, the Phytates are released into ecosystem through dung and due to microbial degradation the phytates are broken down into Phosphates. Phosphate abundance in water bodies leads to Eutrophication. Also many industries inject genetically engineered peptides and steroid hormones to cattle for the improved activity and productivity. Progesterone, Oestradiol, Testosterone and Melengestrol hormones are used mainly in cattle dairies to improve productivity in cattle. These hormones are approved by Food and Drug Administration (FDA) for commercial purpose. Besides productivity, they have certain drawbacks. They cause premature puberty and cancer in consumers (21). Another hormone r-BGH (recombinant Bovine Growth Hormone), is synthetic and used widely in cattle dairies for improving milk productivity. But r-BGH increases the level of IGF-1 (Insulin like Growth factor-1) in milk which can progress various cancers and increases metabolic stress in the body (22).

Another ecological issue that is bothering is stubble burning in many parts of India. Stubble burning is an intentional burning of post-harvest paddy/wheat straw unused. This major causes air pollution. The practice of stubble burning raises the concentration of particulate matter (PM) in air triggering the medical emergency cases. Recent satellite studies indicated the movement of smoke from the regions of Haryana, Punjab and towards Delhi. These contributions magnify the air pollution in Delhi (23).

Paddy straw is the conventional source of animal fodder to ruminant animals. It is rich in Cellulose, Hemicellulose and lignin contents i.e., Cellulose (32-47 %), Hemicellulose (19-27%) and Lignin (5-24%) (24, 25). It promotes fat content in the milk. Here, we used paddy straw as the major component of the feed. Azolla is cultivated, dehydrated and then powdered. Hay powder and Azolla powder are mixed along with jaggery as binder. Jaggery adds taste to the feed and attracts cattle to feed them. Also jaggery acts as binder for easy pelleting. Since Azolla is rich in dietary fiber and crude proteins, it is chosen as one of the ingredient in NuCa feed. Azolla improves the milk yield by 10-12 % in cattle (13, 14). These feed pellets are early in packing and transporting. They are resistant to adulteration and spoilage. Azolla cultivation concurrently produces biomanure After cultivating the Azolla in bulk, spent slurry in the tank can be used as biofertilizer for crop improvement. This biomanure has low C/N ratio i.e., high Nitrogen and low Carbon content. The current feed products mainly comprise cornmeal, soya meal and others which are also food crops for humans and they contain Phytate which is an anti-nutritional factor and they take away significant amount of cultivated land. To overcome feed shortage many industries inject genetically engineered hormones to improve the milk yiled in cattle. The main problem associated with these hormones is disturbance in body metabolism, cancer progression and premature puberty in children before reaching adolescent age. Feed developed in this study is free from phytate and does not encroach into human food share and farmland that is meant for food crops. In fact, it is prepared from the agricultural residue i.e., paddy straw and Azolla. In many parts of the country, the crop straw and stubble are burnt for lack of technology to store it for cattle feed for a long period. Though it is established that paddy straw improves milk quality (by increasing the fat content in milk), there is no technology at present in market to produce paddy straw as a viable product available throughout the year in all the places. Concurrently, Azolla is a good source of minerals and proteins which can be easily grown in backyards. The backyard Azolla cultivation in the rural and urban areas enriches the women rural empowerment. This will be an add on income generation for the household women in the rural and urban sectors. Protein deficiency is one of the major requirment for both livestock and the human population nowadays, this feed improves productivity, to fight against protein hunger and improve the nutritional standards. A combination of Hay and Azolla can provide necessary nutrients for improved milk yield and quality in cattle. A product from Hay & Azolla can replace the existing feed which has several setbacks. Another competitive advantage is an increase in milk production and the price is much cheaper than the existing feeds. This study enabled us to understand the problems in livestock sector due to feed shortage and importance of Azolla as animal feed. NuCa feed is produced by combining Azolla and paddy hay powder as major components. The process for feed preparation is fast and cheap in cost than existing commercial feeds. And also rural farmers can produce feed by their own with this technology and the method that we employed here for azolla is very cheaper than existing methods.

## Acknowledgement

We thank Vignan’s Foundation for Science Technology and Research, Vadlamudi, Guntur, AP for providing in-house fund for procuring machinery and materials in carrying out the research work and Head of the Department, Biotechnology for the strong support.

